# popDMS infers mutation effects from deep mutational scanning data

**DOI:** 10.1101/2024.01.29.577759

**Authors:** Zhenchen Hong, John P. Barton

## Abstract

Deep mutational scanning (DMS) experiments provide a powerful method to measure the functional effects of genetic mutations at massive scales. However, the data generated from these experiments can be difficult to analyze, with significant variation between experimental replicates. To overcome this challenge, we developed popDMS, a computational method based on population genetics theory, to infer the functional effects of mutations from DMS data. Through extensive tests, we found that the functional effects of single mutations and epistasis inferred by popDMS are highly consistent across replicates, comparing favorably with existing methods. Our approach is flexible and can be widely applied to DMS data that includes multiple time points, multiple replicates, and different experimental conditions.

---

Understanding the relationship between protein sequence and phenotype is a central question in evolution and protein engineering. In recent years, a new family of experimental methods, commonly referred to as deep mutational scanning (DMS) or multiplexed assays for variant effects (MAVEs), have been developed to measure the functional effects of large numbers of mutations simultaneously ^1,2^. DMS experiments typically work by generating a vast library of protein variants that are then passed through rounds of selection that favor functional variants while eliminating inactive ones ^3^. One can then compare variant frequencies in the pre- and post-selection libraries to estimate the functional effects of mutations. This approach has been successfully applied in a wide variety of contexts, from studying the function of enzymes ^4^ and tRNAs ^5^ to measuring the mutational tolerance of influenza ^6–8^ and human immunodeficiency virus (HIV-1) ^9–11^ surface proteins.

Despite the success of DMS experiments, popular approaches for analyzing DMS data yield surprisingly modest correlations between the inferred functional effects of mutations in experimental replicates. Thus, a significant amount of variance in the data remains unexplained. Some methods use the ratios between post- and pre-selection variant frequencies, known as enrichment ratios, to estimate mutation effects ^12–14^. Ratio-based methods may be sensitive to noise when variant counts are low, a common occurrence in DMS experiments. Methods based on regression ^15–19^ provide improved performance, but substantial uncertainty in the inferred effects of different mutations persists.

We developed a method, popDMS, to estimate the functional effects of mutations in DMS experiments using statistical methods from population genetics (Methods). In our approach, we view rounds of phenotypic selection in experiments as analogous to rounds of reproduction in natural populations. We quantify the effect of each mutation *i* by a selection coefficient *s*_*i*_, which describes the relative advantage or disadvantage of the mutation for surviving selection in the experiment. Leveraging recently-developed computational methods ^20–22^, we can quantify the likelihood of experimentally observed variant frequency changes as a function of the selection coefficients. We then use Bayesian inference to identify the selection coefficients that best explain the data.

popDMS has several computational strengths. First, the use of a prior distribution for the selection coefficients curbs the inference of strong functional effects in the absence of strong statistical evidence. Our Bayesian framework further allows us to derive joint estimates of selection coefficients across replicates that are guided by levels of evidence in the data, rather than simply averaging the inferred functional effects of mutations across replicates. When information about sequencing error rates is available, we can perform error correction for variant frequencies. In simulations, we found that popDMS was robust to sampling noise and provided stronger correlations between inferred variant effects across replicates than common methods based on enrichment ratios or regression (**Supplementary Fig. 1**).

Next, we analyzed a collection of 25 DMS data sets with popDMS ^5,11,15,16,23–31^. These data sets were generated and analyzed using a variety of experimental techniques and analytical methods (see **Supplementary Table 1**). Like the functional metrics introduced by previous methods, selection coefficients provide an intuitive visualization of the functional effects of mutations (**Fig. 1a**). To quantify the consistency of different analytical methods, we computed the Pearson correlation *R* between mutation effects inferred from replicates of the same experiment. We found that mutation effects inferred by popDMS had higher correlations between replicates than those inferred by prior methods for all the data sets that we considered (**Fig. 1b**). Our selection coefficients also compared favorably with the frequencies of amino acid variants in influenza viruses in a natural population ^6^ (see Methods). To illustrate performance in a typical case, we show selection coefficients inferred for mutations in the HIV-1 envelope protein BF520 (**Fig. 1c**) compared with enrichment ratios (**Fig. 1d**) for the same data ^11^. Improvements in consistency across replicates with popDMS were often substantial. The mean improvement in *R*^2^ for variant effects was 0.36, with 6 out of 25 data sets showing an improvement in *R*^2^ of *>*0.50 (**Fig. 1e**).

**Fig. 1.**
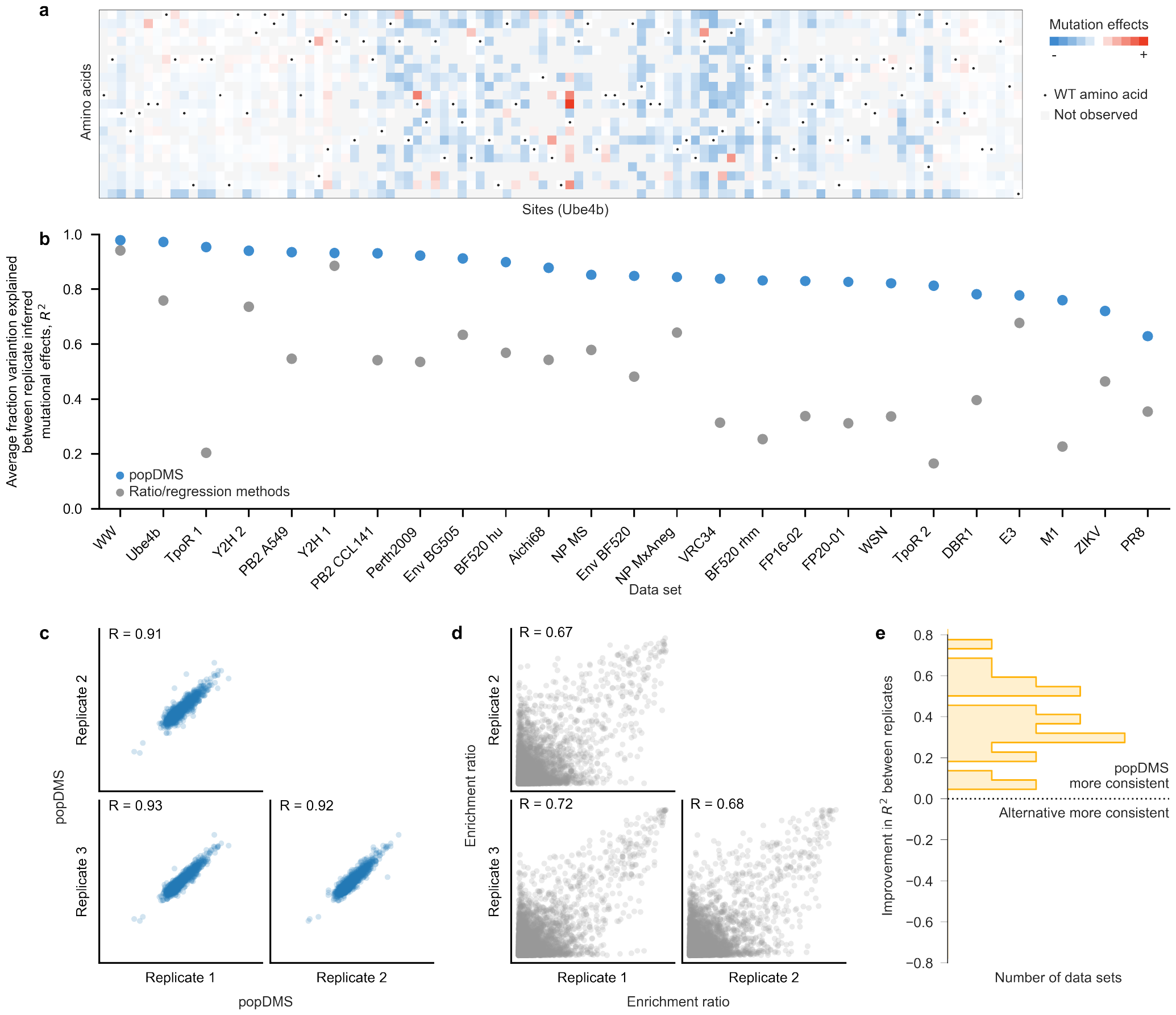
popDMS overview. **a**, Example of the effects of mutations inferred by popDMS for the Ube4b protein ^23^. **b**, Across 25 data sets, popDMS infers more consistent mutational effects than previous ratio/regression-based methods. To illustrate consistency between replicates, we show (**c**) selection coefficients inferred across replicates for the HIV-1 envelop BF520 data set ^11^, compared with (**d**) enrichment ratios for the same data. **e**, popDMS gains in consistency across replicates are often substantial, improving *R*^2^ by an average of 0.36.

We then asked how similar the selection coefficients inferred by popDMS are to mutation effects inferred by previous methods. Across the experimental data sets that we tested, popDMS results were broadly consistent with existing metrics (average Pearson’s *R* = 0.70). This correlation is similar to the average correlation between replicates of the same data set using current ratio- or regression-based methods (average Pearson’s *R* = 0.69). **Figure 2a** shows a typical example, comparing selection coefficients inferred by popDMS with enrichment ratios for the HIV-1 Env BG505 data set ^31^. While the inferred mutation effects agreed for most sites, some showed qualitative differences (**Supplementary Fig. 2**). One factor underlying this result is that popDMS models variants with high initial frequencies, such as wild-type or reference amino acids, in the same way as other, low-frequency variants (see Methods). In alternative methods, the statistical treatment for wild-type amino acids is often different than for other variants.

**Fig. 2.**
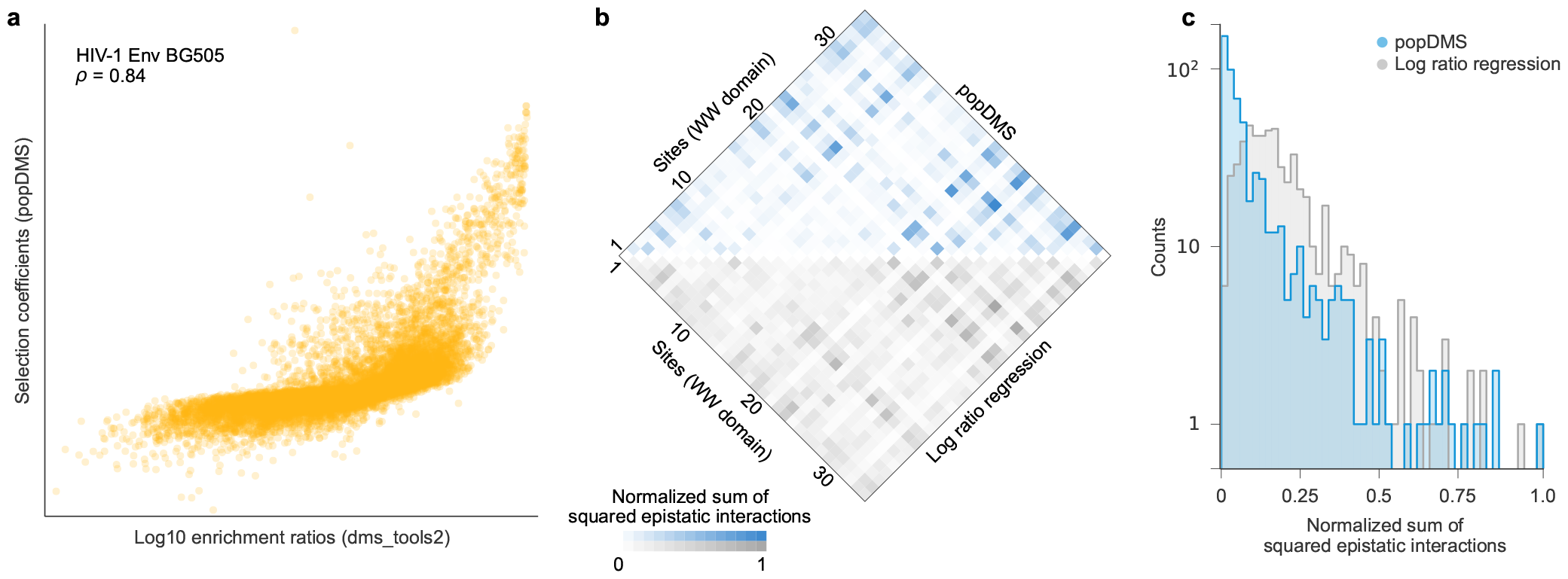
Mutation effects inferred by popDMS are broadly consistent with alternative methods. **a**, For the HIV-1 Env BG505 data set, selection coefficients inferred by popDMS are congruent with enrichment ratios computed using dms_tools2 (Spearman’s *ρ* = 0.84). At some sites, significant differences are observed (see **Supplementary Fig. 2**). **b**, In the hYAP65 WW domain data set, similar sites are inferred to have strong epistatic interactions using popDMS and log ratio regression ^15^. Interactions inferred in ref. ^15^ have been transformed to compare more directly with interactions inferred by popDMS, and both sets of interactions are normalized to scale between zero and one (Methods). **c**, Epistatic interactions inferred by popDMS are substantially sparser than those inferred with the regression-based approach ^15^.

Beyond inferring the effects of individual mutations, we can apply popDMS to estimate pairwise epistatic interactions between variants at different sites. We inferred epistatic interactions in an hYAP65 WW domain data set using popDMS, which we also compared with previous results ^15^. Due to different conventions in defining epistasis, we transformed the functional measurements defined in ref. ^15^ to more directly compare with our results (Methods). To more clearly identify strongly interacting pairs of sites, we computed the sum of squared epistatic interactions between all pairs of amino acids at each pair of sites in the WW domain, using both popDMS and the previous regression-based approach. Our results showed good agreement with the pairs of sites that were previously inferred to have the strongest epistatic interactions (**Fig. 2b**). However, epistatic interactions inferred by popDMS were substantially sparser than those that had been inferred before (**Fig. 2c**). Given the enormous number of possible epistatic interactions between amino acid variants at different sites, sparsity is an attractive statistical feature that can facilitate focus on a smaller number of biologically important interactions.

In summary, popDMS is an efficient, reliable approach for inferring mutation effects from DMS data, which is grounded in evolutionary theory. Across simulations and a wide array of data sets, we found that popDMS infers more consistent mutation effects than the popular alternatives used here. Our approach allows us to combine statistical power across multiple replicates, and it is also capable of inferring epistatic interactions given appropriate data. popDMS is written in Python3 and C++, and uses codon counts in dms_tools format ^14^ or sequence counts in MaveDB format ^32^ as input, with code and example visualizations freely available on GitHub (https://github.com/bartonlab/popDMS, Methods).

## ACKNOWLEDGEMENTS

The work of Z.H. and J.P.B. reported in this publication was supported by the National Institute of General Medical Sciences of the National Institutes of Health under Award Number R35GM138233.

## AUTHOR CONTRIBUTIONS

All authors contributed to methods development, data analysis, interpretation of results, and writing the paper. J.P.B. supervised the project.

## Methods

### Evolutionary model

We model rounds of selection in deep mutational scanning experiments like rounds of reproduction in an evolving population. For this purpose, we use the Wright-Fisher (WF) model ^33^, a simple model from population genetics where individuals in a population undergo discrete rounds of mutation, selection, and reproduction. We define the Wright-Fisher model as follows. We assume that the population consists of *N* individuals, each of which possesses a genetic sequence of length *L*. Each site in the genetic sequence can take on one of *q* possible states, resulting in *M* = *q*^*L*^ possible genotypes.

In the context of DMS experiments, we are typically interested in the properties of proteins with different amino acid variants at each site, and thus we use *q* = 21 for data analyses (representing 20 amino acids and a stop, which could also be further extended to account for gaps). However, the framework that we consider is more general. One could consider nucleotide sequences with *q* = 4 states (A, C, T, G), *q* = 64 codons, and so forth.

At each time *t*, the state of the population is defined by a genotype frequency vector ***z***(*t*) = (*z*_1_(*t*), *z*_2_(*t*),…, *z*_*M*_ (*t*)), where *z*_*a*_(*t*) = *n*_*a*_(*t*)*/N*, with *n*_*a*_(*t*) representing the number of individuals that have genotype *a* at time *t*. Under the WF model, the probability of observing genotype frequencies ***z***(*t* + 1) in the next generation is binomial,

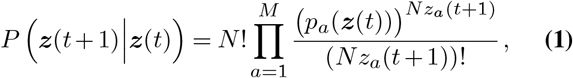

with

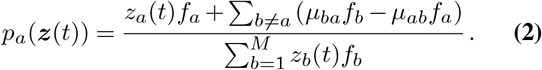

Here *f*_*a*_ is the *fitness* of genotype *a*, defined in detail below, and *µ*_*ab*_ is the probability of mutation from genotype *a* to genotype *b* in one generation. In typical experiments, mutation rates are low enough that we assume *µ*_*ab*_ is zero across all pairs of genotypes *a, b*.

We assume that the fitness of each genotype depends linearly on the amino acid (or nucleotide, codon, etc.) content of the sequence,

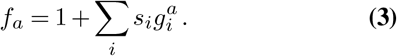

In Eq. (3), the *s*_*i*_ are *selection coefficients* for each variant *i*, which quantify the effect of that variant on fitness. If *s*_*i*_ is positive, then the variant is beneficial, and if *s*_*i*_ is negative, then the variant is deleterious. Here 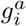 is an indicator variable, which is equal to one if genotype *a* possesses the variant *i*, and zero otherwise. The variant indicator *i* is a generic index that runs across all possible amino acids or states at each site in the sequence. For example, let us define a genotype sequence *a* = *{T, E, K}*. For this sequence, 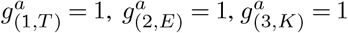 and all other 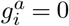.

Following Eq. (1), the probability of a sequence of *K* genotype frequency vectors *{ z*(*t*_1_), *z*(*t*_2_), …, *z*(*t*_*K*_) *}*, conditioned on an initial distribution of genotype frequencies *z*(*t*_0_), is given by the product of the individual transition probabilities,

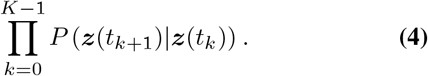

### Inferring fitness effects of mutations with popDMS

We view sequencing results in a DMS experiment as measurements of the genotype frequency vectors *z*(*t*). To infer the functional effects of mutations, we take a Bayesian approach, seeking the selection coefficients *s* that maximize the posterior probability of the entire evolutionary trajectory Eq. (4). This includes a Gaussian prior distribution for the selection coefficients

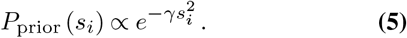

Here *γ* encodes of the width of the prior distribution, which can also be thought of as *L*_2_-norm regularization of the selection coefficients. The overall posterior distribution for the selection coefficients is then given by

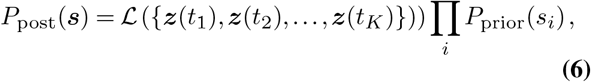

where the likelihood of the data *ℒ* is given by Eq. (4).

Following recent computational advances ^34^, to simplify the likelihood, we consider the diffusion limit of the WF model. In this limit, we assume *N* is large and the *s* and *µ*_*ab*_ are small. This transforms Eq. (4) from a complicated function of the selection coefficients into a Gaussian, though Eq. (4) retains a complex dependence on the genotype frequencies. The maximum *a posteriori* (MAP) estimate for the selection coefficients *ŝ* is given by

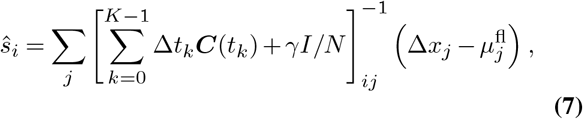

where Δ*t*_*k*_ = *t*_*k*+1_ *− t*_*k*_, Δ*x*_*j*_ = *x*_*j*_(*t*_*K*_) *− x*_*j*_(*t*_0_), and 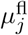 is the net expected change in the frequency of variant *j* over the course of the experiment due to mutation alone. Typically, *µ*_fl_ is assumed to be zero, except for experiments involving viral replication, where mutation rates can be high enough to produce observable changes in frequency. Here *C*(*t*) is the covariance matrix of variant frequencies 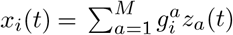 which has entries

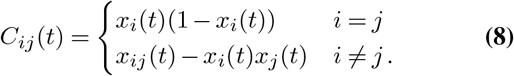

Here 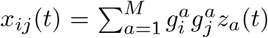 is the frequency of genotypes at time *t* that contain both variants *i* and *j*.

The estimate of the selection coefficients *ŝ* given in Eq. (7) can be explained intuitively. First, for simplicity, consider the matrices *C*(*t*_*k*_) to be diagonal. Then, the estimate for *ŝ* depends on how much variant *i* has increased in frequency over the course of the experiment, after correcting for changes in frequency that are not due to functional selection, 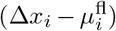 This quantity is normalized by the variance of the variant frequency *x*_*i*_(*t*_*k*_) over time (Eq. (8)). In the limit that *x*_*i*_(*t*_*k*_) is small (and again, that the off-diagonal terms are zero), the estimate for *ŝ* _*i*_ is similar to an enrichment ratio, because in this limit 1 *− x*_*i*_(*t*_*k*_) *≈* 1. However, this estimate is also shrunk by a factor of *γ* due to the prior distribution for the selection coefficients. Importantly, the variance also becomes small when *x*_*i*_(*t*_*k*_) is close to one, as is often the case for wildtype (WT) or reference amino acids in DMS experiments. This distinguishes the treatment of WT variants in popDMS as compared to ratio-based methods and regression-based methods that do not assume logistic growth.

Off-diagonal terms in Eq. (7) account for the influence of genetic background on changes in variant frequency. For example, a variant *i* may increase in frequency not because it has a beneficial functional effect, but rather because it appears on the same genetic sequence with other beneficial variants more often than expected by chance (i.e., positively covarying with other beneficial variants; see Eq. (8)). In population genetics, this phenomenon is referred to as genetic hitchhiking ^35^. In DMS data, covariances cannot always be computed due to limited read lengths, but this information can be used to enhance predictions when it is available.

To derive Eq. (7), we assumed that the number of individuals in the population, *N*, is constant. However, in experiments (and in real populations), *N* can vary in time. Incorporating time-varying population sizes leads to similar estimates of selection, but with a larger uncertainty in the inferred parameters (see ref. ^36^ for a related model). For simplicity, we will maintain the assumption that *N* is constant. Additionally, in the discussion below we will absorb the population size *N* into the definition of *γ*, so that the strength of the prior distribution does not rely on an arbitrary definition of population size.

### Joint estimates of selection coefficients across experimental replicates

We model experimental replicates as alternative evolutionary histories, subject to the same functional effects of mutations but with different stochastic realizations of evolution (and potentially different starting conditions). The posterior probability for the selection coefficients across *R* replicates is then given by

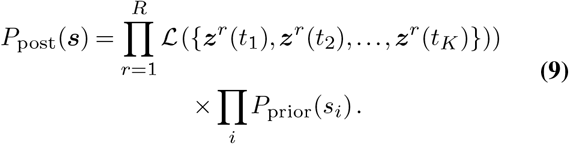

Here each experimental replicate has a different index *r*, and the likelihood across all replicates is the product of the likelihood for each individual replicate. Since each *ℒ* is Gaussian in the selection coefficients, the product is also Gaussian, and the MAP selection coefficients can be computed as in Eq. (7), yielding

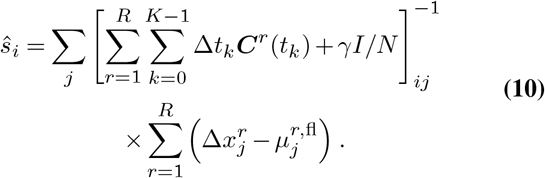

### Correction for sequencing errors

For some data sets, information on sequencing error rates is available. For example, this can be obtained by sequencing a library consisting of all WT sequences, so that all differences from WT are likely attributable to sequencing errors. When this data is available, we compute corrected mutant and WT counts by subtracting the expected contributions from sequencing errors.

### Optimizing the regularization strength

For simplicity, we incorporate the WF population size *N* into the prior parameter *γ* to define an effective regularization strength *γ*^*′*^ = *γ/N*. Larger values of *γ*^*′*^ put a higher penalty on inferred selection coefficients, thereby suppressing their values, but also limiting the effects of sampling noise. Smaller values of *γ*^*′*^ allow for the inference of larger selection coefficients, but in turn, these estimates are more sensitive to noise.

One can choose a single value of *γ*^*′*^ to use for all data sets, but this parameter can also easily be optimized for an individual data set. The most computationally intensive step in inferring mutation effects (i.e., selection coefficients) with popDMS is computing the variant frequencies and covariances from sequencing data. After this step has been completed, it is straightforward to sweep through a range of *γ*^*′*^ values and test their results for each data set.

We found that the average correlation of inferred mutation effects between replicates typically behaves like a logistic function of log (*γ*^*′*^). For very small values of *γ*^*′*^, sampling noise is not effectively suppressed, and the correlation of inferred mutation effects between replicates is lower. As *γ*^*′*^ increases, noise is suppressed, leading to higher correlations between replicates. At high values of *γ*^*′*^, high correlations between replicates are typically preserved, but the inferred selection coefficients are shrunk strongly towards zero.

We reasoned that an optimal choice for the regularization strength *γ*^*′*^ would be the smallest value of *γ*^*′*^ that effectively suppresses sampling noise, as this would avoid shrinking estimated selection coefficients unnecessarily. To compute this value, for each experimental data set described below, we swept through values of *γ*^*′*^ in even logarithmically spaced steps from roughly 1*/B*, where *B* is the maximum read depth, to 1000. For each value of *γ*^*′*^, we computed the correlation between replicates. We then computed the difference Δ*R* = *R*_max_ *− R*_min_ between the maximum correlation and minimum correlation between replicates across all values of *γ*^*′*^. To determine the optimum value of *γ*^*′*^, we started with the value that corresponds to the maximum correlation between replicates and moved to progressively smaller values, stopping when the correlation drops by more than 10% of Δ*R* from one value of *γ*^*′*^ to the next.

While sweeping through values of *γ*^*′*^ improves our consistency across data sets, allowing us to adjust our regularization to match the level of noise in the data, we emphasize that this step is not essential to obtain robust results. A simple choice of *γ*^*′*^ = 0.1 is nearly optimal for every data set we considered, with the exception of the influenza PR8 study ^37^. This data set is the only one in which the correlation between replicates is not roughly a logistic function of the regularization strength.

### Generating logo plots with popDMS

Inferences from DMS data such as amino acid preferences (derived from enrichment ratios) have often been used to generate logo plots that show the relative dominance of different amino acids at each site. However, while preferences naturally sum to one, selection coefficients inferred by popDMS can be both positive and negative. To obtain preference-like logo plots using selection coefficients inferred by popDMS, computed exponentially transformed values

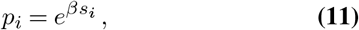

where the scaling factor *β* was approximately chosen to maximize the correlation between the transformed selection coefficients *p*_*i*_ and amino acid preferences for the same data set.

### Inference of epistasis

We extended our approach to infer pairwise epistatic interactions between variants by adding epistatic interactions *s*_*ij*_ to the previous fitness function Eq. (3), i.e.,

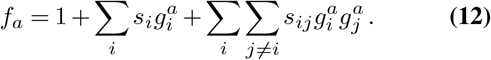

As for the selection coefficients defined above, if an epistatic interaction *s*_*ij*_ is positive, then the presence of variants *i* and *j* together increases fitness more than would be expected from the combined effect of the individual variants. When *s*_*ij*_ is negative, variants *i* and *j* together are more deleterious than expected if they were independent.

With this extension of the fitness model, one can then compute the posterior probability for the change in genotype frequencies, as in Eq. (6). We also assume a Gaussian prior distribution for the epistatic interactions that is centered at zero and has the same width as for the selection coefficients. The MAP selection coefficients for the selection coefficients and epistatic interactions have a form analogous to Eq. (7), but with an expanded index that runs over all variants *i* and all *pairs* of variants (*i, j*). Additional terms in the covariance matrix are then given by

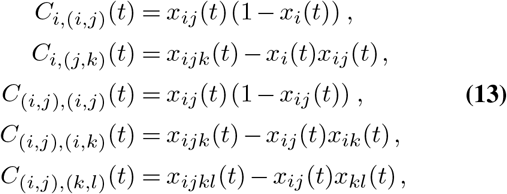

with

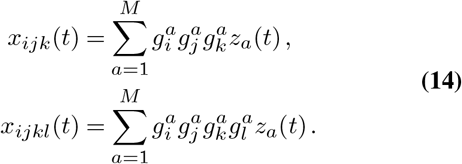

popDMS differs from some alternatives to estimating epistasis in that information about pairwise interactions is gained from all sequences that bear two or more non-reference variants. For example, one previously developed approach effectively estimated the fitness of sequences with exactly two mutations and compared this with estimates of the fitness for corresponding single mutants to estimate the strength of epistatic interaction between the mutations ^38^.

At present, inferring epistatic interactions from DMS data with popDMS is only computationally feasible for short sequences due to the large size of the covariance matrix. Alternative approaches that strictly enforce sparsity and reduce the number of possible interactions to estimate could potentially ease these computational restrictions.

### Testing performance in simulations

We simulated evolution following the WF model over a number of generations to test the performance of popDMS. To reproduce finite sampling statistics similar to those observed in experimental data, we used the initial genotype frequency data from an experimental data set ^39^. We ordered the variants by frequency at each site and inferred a best-fit multinomial model describing the frequency distribution across sites using PyStan ^40^. This inferred distribution thus captures a typical hierarchy of frequencies observed in DMS experiments, from high frequency (WT/reference) variants to rare ones, whose counts may be of the same order as the read depth.

In our simulations, selection coefficients for all variants were chosen at random from a normal distribution with mean zero and standard deviation 0.1. True starting frequencies were sampled at random from the inferred multinomial distribution using PyStan. We then simulated up to 10 generations of evolution following the WF model, here assuming a mutation rate of zero and population size of *N* = 10^8^. From these true trajectories, we obtained finitely sampled frequency trajectories by multinomial sampling from the true frequencies at each generation, with various choices for the sampling depth. To highlight stochasticity, we used a sampling depth of *B* = 5 *×* 10^4^ sequences in **Supplementary Fig. 1a**.

We used this data to compute the average correlation for selection coefficients inferred from different replicates using popDMS, which varies depending on the number of generations of data used (**Supplementary Fig. 1b**). Intuitively, observing the evolution for a longer time leads to more precise estimates.

We compared the results of popDMS against other common approaches, which we implemented as described below. To compute enrichment ratios, we compare the fraction of reads with a particular variant *i* pre- and post-selection,

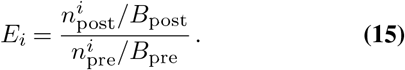

Here 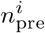 and 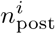 are number of reads with variant *i* before selection and after selection, respectively. Similarly, *B*_pre_ and *B*_post_ represent the total number of reads before and after selection. To compute log ratio scores, we used the natural logarithm of the enrichment ratios,

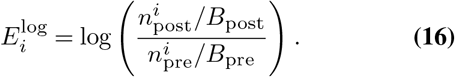

Finally, log ratio regression scores were computed by calculating the logarithm of the enrichment ratio Eq. (16) for each variant at each generation, then extracting the slope of the linear model the best fits the change in log enrichment ratios over time.

### DMS data sets

Data sets used in this paper were obtained from 15 publications ^37–39,41–52^. Additional information about these data sets, and the methods used to analyze them, is summarized in **Table 1**.

### Comparison with prior studies of epistasis

Here we analyzed a data set from Araya and collaborators, which explored epistasis in the WW domain of the hYAP65 protein ^38^. There, they define epistasis in a way that differs from our definition (i.e., the *s*_*ij*_ in Eq. (12)). For each genotype variant *a*, Araya et al. define a parameter 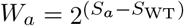, where the *S*_*a*_ are best-fit slopes of the logarithmic enrichment ratios for variant *a. S*_WT_ is the slope for the WT variant, which they use to normalize the results. They use the quantity *ϵ*_*ab*_ = *W*_*ab*_ *− W*_*a*_*W*_*b*_ as the primary metric of epistasis. Here, *a* and *b* represent genotypes with a single mutation, and *ab* the genotype that features only these two mutations.

When the frequency of a variant is small, the *S*_*a*_ computed by Araya et al. are similar to our *f*_*a*_. Thus, to compare the quantities inferred by Araya et al. to our *s*_*ij*_, we computed a set of transformed scores, which we write as

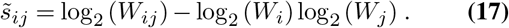

There is good overall agreement in the epistatic interactions *s*_*ij*_ inferred by popDMS and the transformed interactions 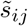, computed from the *W* values of Araya et al. (Pearson’s *R* = 0.73, Spearman’s *ρ* = 0.75). **Figure 2a** similarly shows broad agreement between the sum of squared epistatic interactions between variants at each pair of sites in the WW domain, though those inferred by popDMS are sparser (**Figure 2b**).

### Comparison with natural frequencies of influenza variants

In general, it is challenging to validate inferences about the fitness or functional effects of amino acid variants inferred from DMS experiments because “ground truth” measurements for these effects do not exist. However, one possible method of validation is to compare the inferred fitness effects of variants to the frequency of mutations observed in natural populations. This approach was explored by Thyagarajan and collaborators in their study of the effects of mutations in the influenza hemagglutinin protein ^48^.

We performed a similar analysis to compare our results to fitness effects inferred using enrichment ratios for the same data set ^48^. While it is possible to directly correlate variant frequency and the inferred fitness effect of the variant, this connection is not entirely natural because frequency should be determined not just by the fitness effect of a variant, but also by the relative fitness effects of other possible variants at the same site.

To make a clearer connection with the data, we reasoned that, in most cases, the amino acid with the highest frequency in natural populations should be the variant with the highest fitness at each site. We thus ranked the fitness effects of each amino acid variant at the same site, and computed the rank of the top variant according to both selection coefficients inferred by popDMS and enrichment ratios. For popDMS, the amino acid most frequently observed in natural populations had an average rank of 2.1 across sites (median 1), compared to an average rank of 2.7 (median 1) for enrichment ratios.

To determine the extent to which the amino acid that is most frequently observed in natural populations is predicted to be dominant at each site, we also computed a *z* score for the most frequent variant at each site. This was computed by taking the metric of fitness (selection coefficients or enrichment ratios) for the most frequent variant at each site, subtracting the mean value for the same site, and dividing by the standard deviation of values at that site. We found an average *z* score for the most frequent variant of 3.5 using popDMS, compared to 2.6 for enrichment ratios.

Thus, we find that selection coefficients match well with the corresponding frequencies of amino acid variants in a natural population. Results obtained using popDMS also compare favorably with prior results computed using enrichment ratios ^48^.

## Data and code

Raw data and code used in our analysis are available in the GitHub repository located at https://github.com/bartonlab/paper-DMS-inference. This repository also contains Jupyter notebooks that can be run to reproduce the results presented here. Code for popDMS alone, without the analysis contained in this paper, is also provided in a separate GitHub repository at https://github.com/bartonlab/popDMS. popDMS is coded in Python3 and C++.

---

**Supplementary Table 1.**
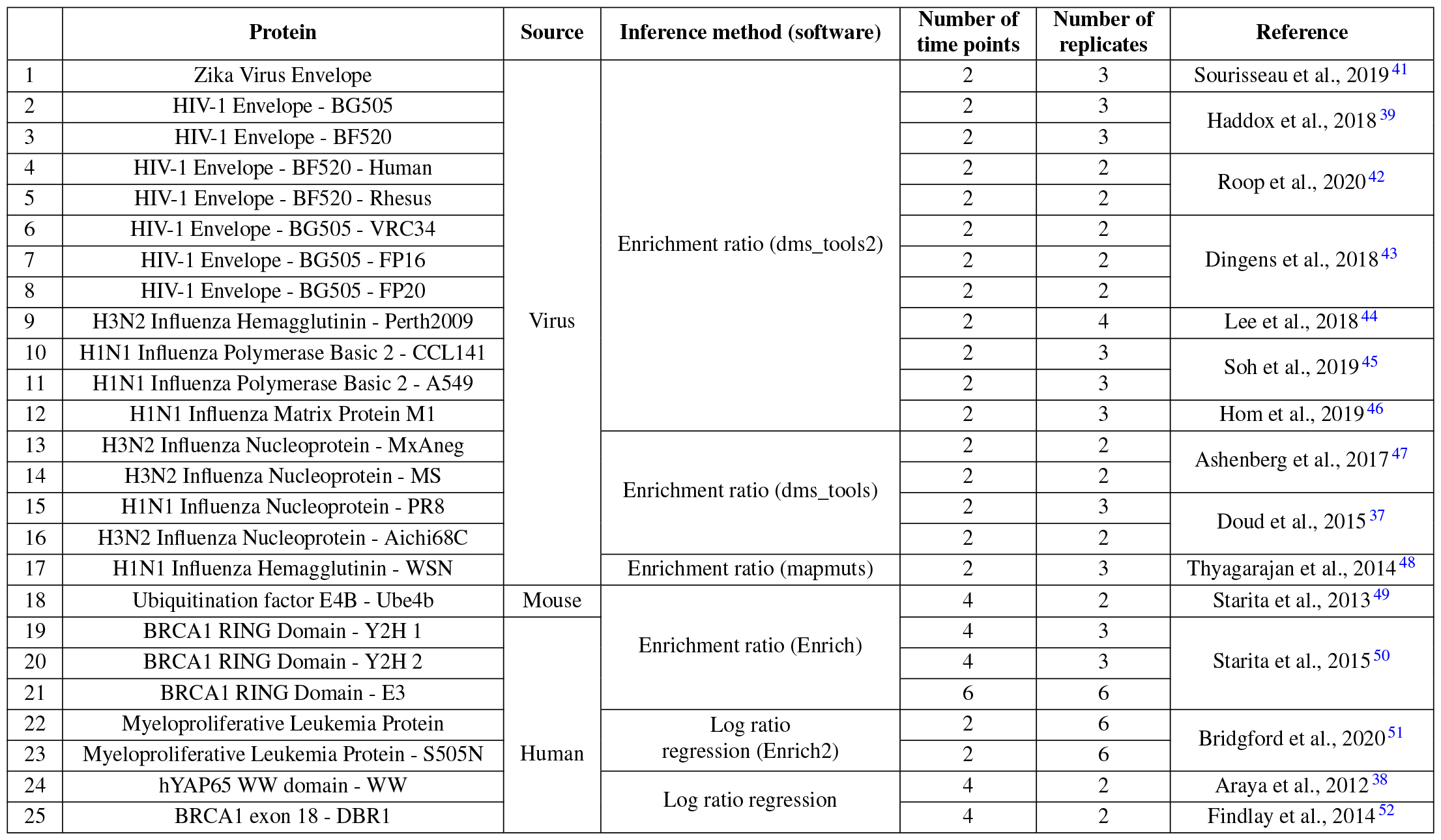
Summary of data sets studied in this work.

**Supplementary Fig. 1.**
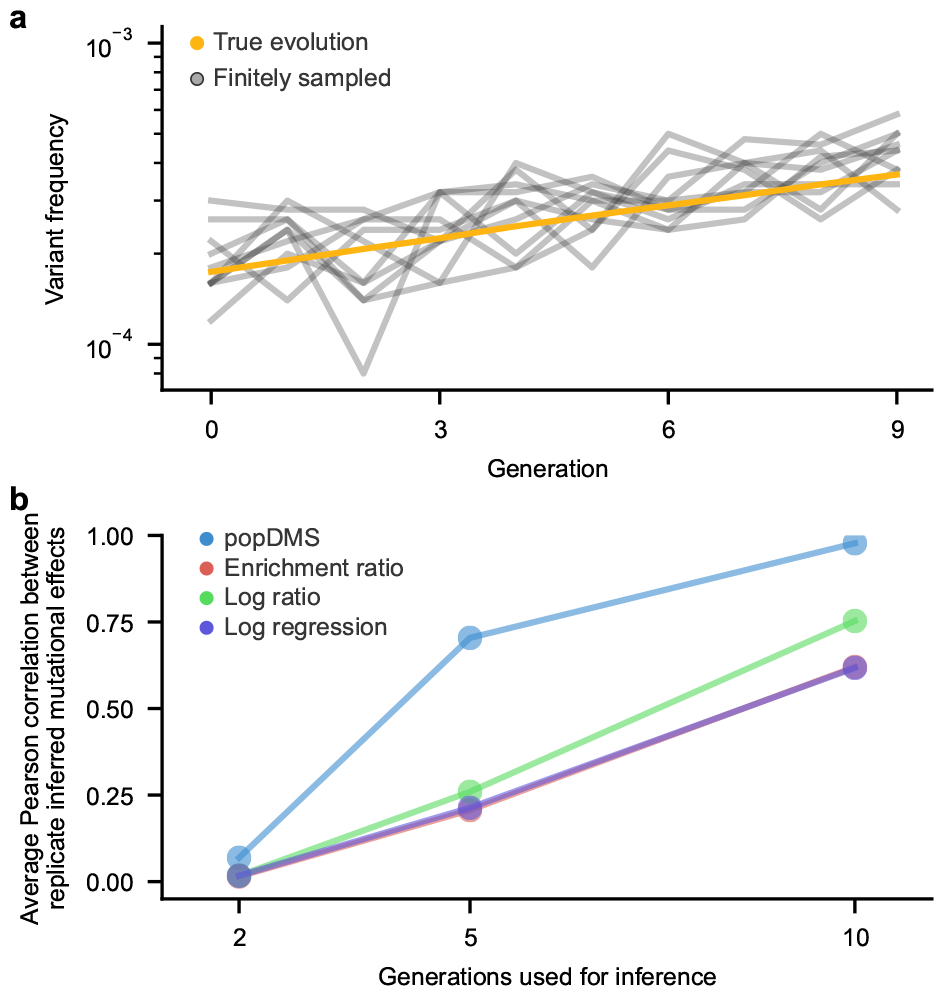
popDMS is robust to finite sampling error. **a**, Due to finite sampling of the data, variant frequencies can appear to fluctuate over time even if the underlying behavior is smooth, complicating inference. Results from an example simulation (Methods). **b**, As the number of generations used for inference in simulations increases, all methods become more robust. popDMS is especially robust in inferring mutation effects from limited data with few rounds of selection.

**Supplementary Fig. 2.**
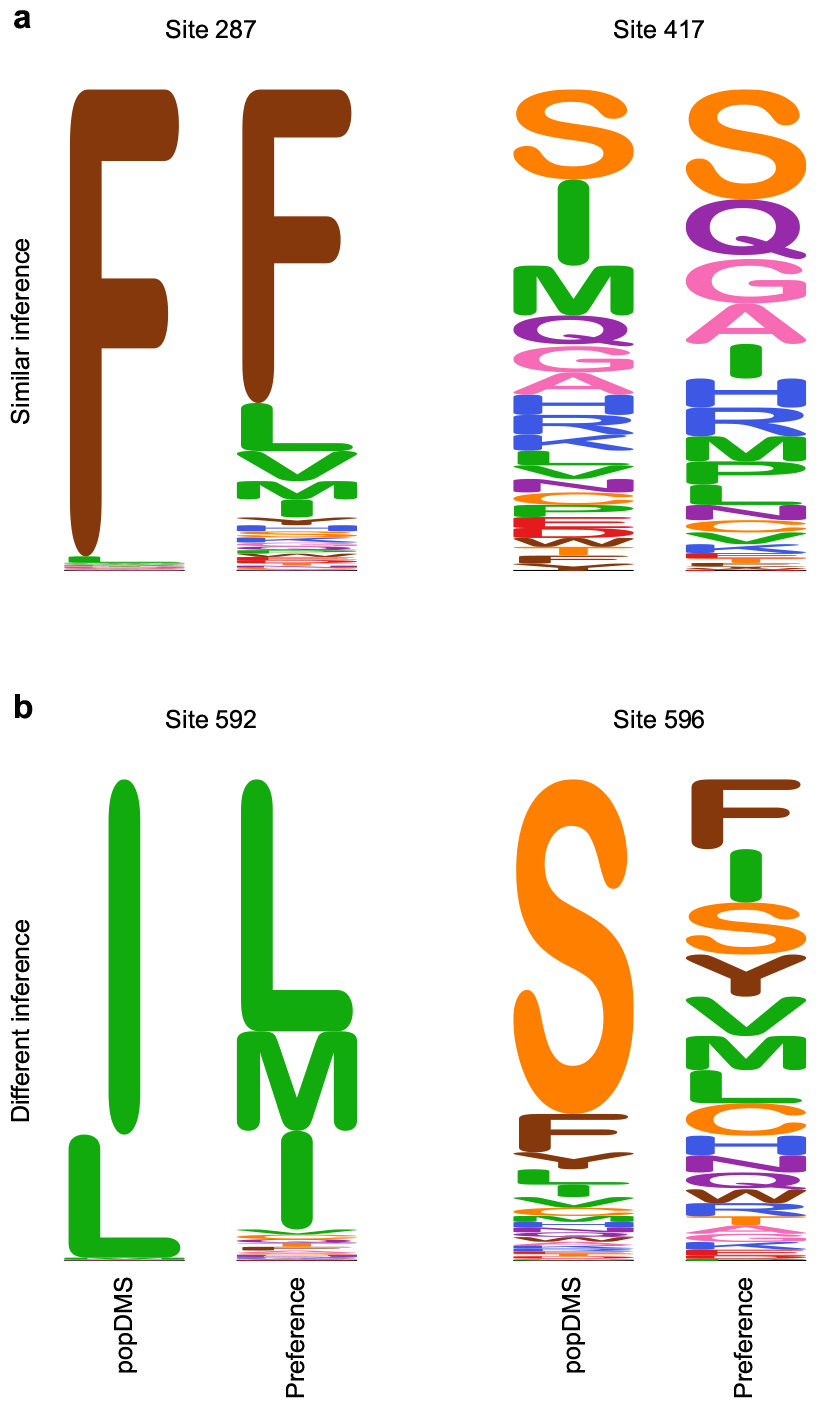
Comparison of selection at individual sites inferred by popDMS and enrichment ratios for the HIV-1 Env BG505 data set. **a**, Exponentially transformed selection coefficients inferred by popDMS (see Methods) are similar to preferences (normalized enrichment ratios) at sites 287 and 417. At site 287, both methods agree on the dominance of phenylalanine. At site 417, both methods find broad tolerance for different amino acid variants. **b**, In contrast, differences are observed between popDMS and preferences at sites 592 and 596. In both cases, popDMS finds the reference amino acid (isoleucine at site 592 and serine at site 596) to be strongly favored due to its increase in frequency during the experiment. These frequency changes were small relative to the initial frequency of the amino acid, but they were large considering the limited capacity for the amino acid to grow in frequency. This latter factor is captured by popDMS, but is not typically accounted for in ratio-based approaches.

